# Polymeric Pathogen-like Particles-Based Combination Adjuvants Elicit Potent Mucosal T Cell Immunity to Influenza A Virus

**DOI:** 10.1101/2020.07.10.197749

**Authors:** Brock Kingstad-Bakke, Randall Toy, Woojong Lee, Pallab Pradhan, Gabriela Vogel, Chandranaik Marinaik, Autumn Larsen, Yoshihoro Kawaoka, Krishnendu Roy, M Suresh

**Author notes:** Corresponding Authors Krishnendu Roy, M. Suresh. Lead Contact : M. Suresh.

## Abstract

Eliciting durable and protective T cell-mediated immunity in the respiratory mucosa remains a significant challenge. Polylactic-co-glycolic acid (PLGA)-based cationic pathogen-like particles (PLPs) loaded with TLR agonists mimic biophysical properties of microbes and hence, simulate pathogen-pattern recognition receptor interactions to safely and effectively stimulate innate immune responses. We generated micro particle PLPs loaded with TLR4 (glucopyranosyl lipid adjuvant, GLA) or TLR9 (CpG) agonists, and formulated them with and without a mucosal delivery enhancing carbomer-based nanoemulsion adjuvant (ADJ). These adjuvants delivered intranasally to mice elicited high numbers of influenza nucleoprotein (NP)-specific CD8+/ CD4+ effector and tissue-resident memory T cells (TRMs) in lungs and airways. PLPs delivering TLR4 versus TLR9 agonists drove phenotypically and functionally distinct populations of effector and memory T cells. While PLPs loaded with CpG or GLA provided immunity, combining the adjuvanticity of PLP-GLA and ADJ synergistically enhanced the development of airway and lung TRMs and protective immunity to pathogenic influenza A virus. Further, balanced CD8 (Tc1/Tc17) and CD4 (Th1/Th17) recall responses were linked to effective influenza virus control in the lungs. These studies provide mechanistic insights into vaccine-induced T cell immunity in the respiratory tract and pave the way for the development of a universal influenza vaccine.

## INTRODUCTION

Respiratory infections in adults and children have been among the top three leading causes of death and disability in the world for decades^1,2^. Novel respiratory pathogens are emerging and can quickly spread due to the ease of transmission, as witnessed in the current coronavirus^3^ and past influenza pandemics^4^. Vaccines are essential for the control and elimination of these diseases by eliciting antibody and/or T cell-mediated immune responses (CMI). The necessity for effective T cell-based vaccines to respiratory pathogens is exemplified by continued seasonal influenza endemics and sporadic pandemics, despite wide vaccine availability and high virus infection rates.

Widely used inactivated influenza vaccines (IIV) function by eliciting subtype-specific antibodies to mutation-prone surface hemagglutinin (HA) and neuraminidase (NA) proteins, and must be reformulated annually to match seasonal strains due to antigenic drift and shift^5^. Further, these IIVs are poorly immunogenic for CD8 T cells. Influenza viral infection can generate strong antibody and T cell-mediated immunity (CMI), and while antibodies are still strongly strain matched in terms of protection, heterosubtypic protection to influenza requires CMI to conserved viral epitopes^5,6^.

Robust T cell control of hetersubtypic influenza infection has been closely associated with the development of mucosally residing tissue resident memory (TRM) CD4+ and CD8+ cells ^7,8^. However, the numbers of TRM cells after influenza infection have been observed to wane over time with a concomitant decrease in heterosubtypic protection^8-10^. Indeed, neither mass vaccination with current IIV or widespread infection have adequately curtailed the continuous infection of influenza virus in human populations. Therefore, development of vaccination strategies to potently elicit long-lived TRM cells and understanding of what factors govern these responses are crucially needed to control respiratory infections such as influenza.

Several groups have developed vaccines to elicit influenza specific lung TRM cells that confer heterosubtypic protection to influenza virus challenge using live attenuated influenza viruses^11^, or viral vectors^12^. Recently, we have tested a novel adjuvant, Adjuplex (ADJ), that when combined with subunit or IIV proteins potently induced CD8+ lung TRM cell responses after mucosal administration, and conferred substantial protection to influenza virus challenge^13^. These studies highlight the importance for identification of novel adjuvants that can elicit mucosal CMI to non-replicating antigens, particularly so we can dissect and study the individual effects these adjuvants have on the magnitude and nature of the resultant mucosal CMI responses.

TLR ligands as adjuvants for influenza vaccines have been widely studied and are typically delivered in monomeric, soluble formulations^14^. In order to mimic biophysical interactions between pattern-recognition receptors and their ligands on pathogens, we have developed TLR agonist-loaded polylactic-co-glycolic acid (PLGA)-based pathogen-like particles (PLPs)^15^. Varying the size of the PLPs and the density of the loaded TLR agonist CpG, modulated the signaling circuitry within dendritic cells *in vitro* and altered the nature of antibody (TH1 versus TH2-driven) responses. Additionally, agonists presented simultaneously on PLPs have been shown to differentially modulate immune responses *in vitro*, compared with soluble counterparts, while potentially improving safety by reducing toxicity from systemic diffusion^16,17^. The ability of TLR-agonist-loaded PLPs to stimulate antigen-specific T cell responses, especially in the respiratory mucosa, has not yet been investigated.

Here, we investigated if CpG- or glucopyranosyl lipid adjuvant^18^ (GLA)-loaded PLP adjuvants mixed with influenza virus nucleoprotein (NP) protein, with or without ADJ could elicit antigen-specific CD4 and CD8 T cells, and analyzed their numbers, frequencies, and phenotypes in systemic and mucosal compartments. We found that PLP vaccine formulations elicited strong, yet phenotypically and functionally distinct TRM CD8+ and CD4+ responses in the lungs of vaccinated mice. Further, we observed that PLP-based adjuvants afforded strong protection from influenza challenge that was closely associated with distinct functional recall profiles of CD4 and CD8 T cells unique to the PLP vaccine formulation. These results highlight that PLP-loaded adjuvants can distinctly program effective CMI and can be leveraged to study immunity and develop vaccines to respiratory pathogens.

## RESULTS

### Adjuplex modifies the response to PLPs with TLR adjuvants in murine BMDCs

We first assessed the extent to which ADJ affected the responses of murine DCs to TLR agonists CpG (TLR9) and monophosphoryl lipid A (MPLA, TLR4) presented in soluble form or as PLPs. Data in Fig. 1 show that both soluble CpG and PLP-CpG triggered potent IFN-β responses from murine BMDCs. Interestingly, ADJ alone did not induce an IFN-β response, but it ablated the IFN-β response induced by both soluble and PLP-CpG (Fig. 1A and 1B). MPLA did not stimulate substantial IFN-β production in either soluble form or as PLP-MPLA (Fig. 1A and 1B). Next, we explored whether GLA (synthetic MPLA) triggered production of pro-inflammatory cytokines such as IL-1β and IL-18 in BMDCs. Neither ADJ nor PLP-GLA alone induced IL-1β or IL-18 production from murine BMDCs, but ADJ + PLP-GLA induced a very strong IL-1β/IL-18 response (Fig. 1C and 1D). Comparison of IL-18 responses to ADJ + PLP-GLA in BMDCs from wild-type, AIM2^-/-^, NLRP3^-/-^, AIM2/NLRP3^-/-^, and ASC^-/-^ mice revealed that the NLRP3 inflammasome is essential for ADJ/GLA-induced IL-1β/IL-18 production by BMDCs (Fig. 1E). Taken together, data in Fig. 1 showed that ADJ in combination with CpG and GLA stimulated disparate cytokine responses from BMDCs.

**Figure 1.**
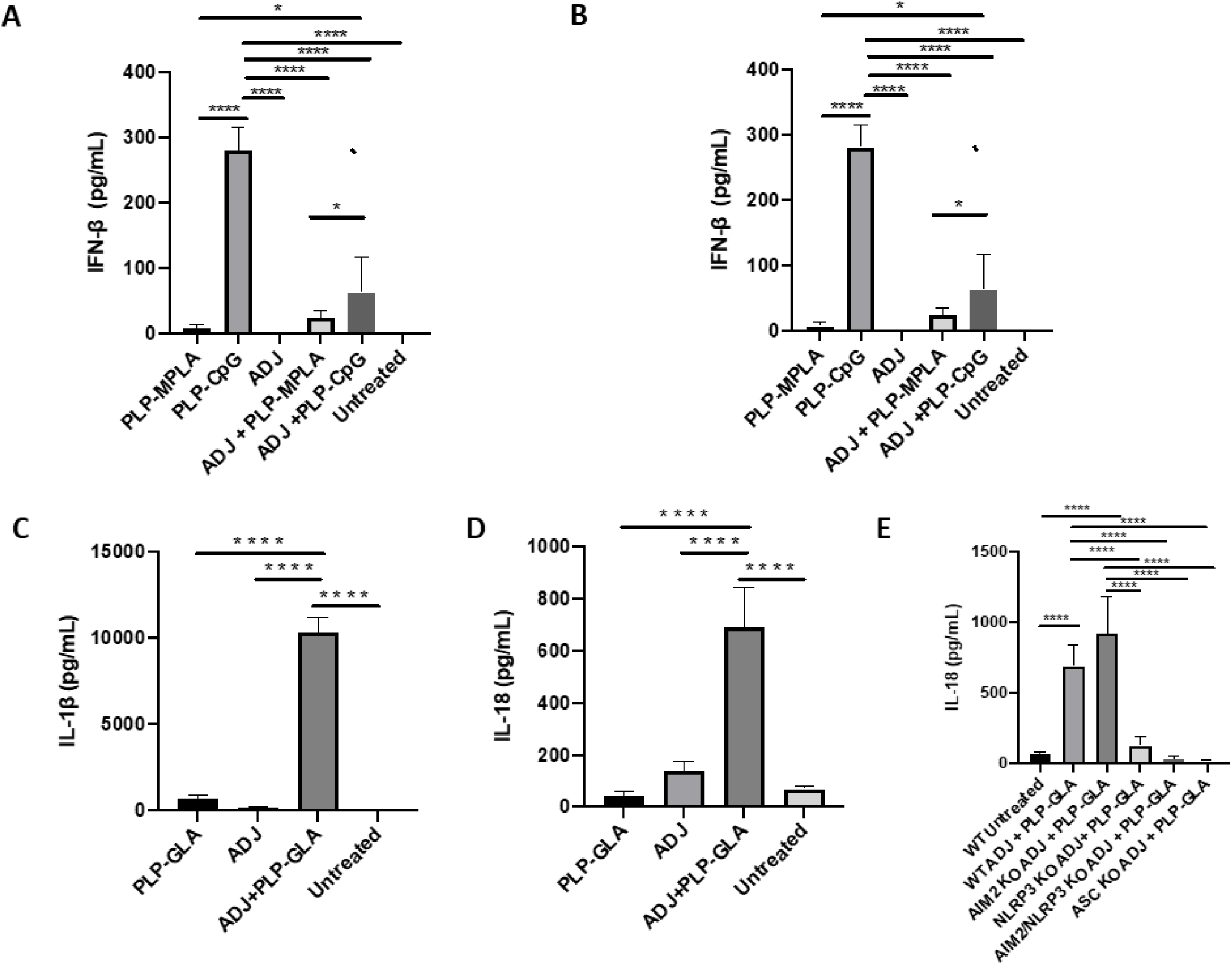
Murine BMDC response to Adjuplex and PLP adjuvants. Murine BMDCs were treated with Adjuplex (ADJ) and/or PLPs with CpG or GLA. (A) IFN-β response from BMDCs after 24 hrs of treatment with ADJ and/or soluble CpG or MPLA. (B) IFN-β response from BMDCs after 24 hrs of treatment with ADJ and/or PLP-CpG or PLP-MPLA. (C) IL-1β and (D) IL-18 response from BMDCs after 24 hrs of treatment with ADJ and/or PLP-GLA. (E) IL-18 response from BMDCs after 24 hours of treatment with ADJ + PLP-GLA in wild-type, AIM2^-/-^, NLRP3^-/-^, AIM2/NLRP3^-/-^, and ASC^-/-^ mice. For A-D, data are representative of two independent experiments. **, ***, and **** indicate significance at P<0.01, 0.001, and 0.0001, respectively.

### Combination PLP adjuvants elicit contrasting CD8 and CD4 T cell effector responses

First, we compared the ability of various TLR agonist-loaded PLPs formulated with or without ADJ to elicit pulmonary CD8 T cell responses to the subunit protein, influenza virus nucleoprotein (NP). All PLP adjuvants elicited robust CD8 T cell responses in the lungs and airways of vaccinated mice (Fig 2A). Remarkably, 30-60% of CD8 T cells in the airways were specific to the immunodominant epitope NP366 and combination with ADJ did not significantly affect the accumulation of such cells in lungs or airways. While PLP-GLA +/- ADJ seemed to drive the highest numbers of NP366-specific CD8 T cells in lungs, PLP-CpG groups had lower levels of these effector cells. Mice administered NP with empty PLP (PLP-E) had relatively low frequencies and numbers of NP366-specific CD8 T cells, indicating that the PLP particles alone were not contributing significantly to induction of this immune response.

**Figure 2.**
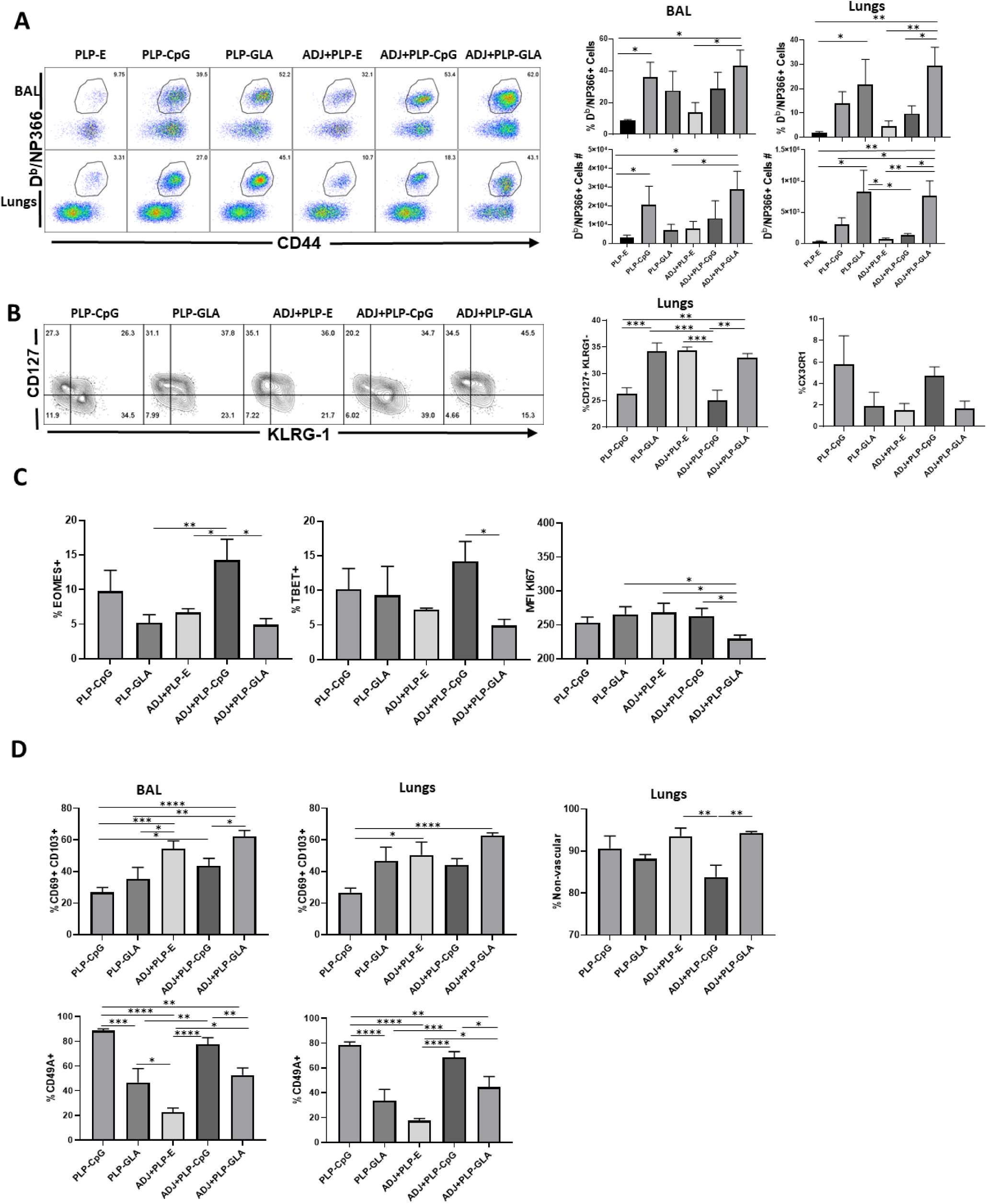
Effector CD8 T cell response to adjuvanted vaccines. B6 mice were vaccinated intranasally twice with influenza virus nucleoprotein (NP) formulated with the indicated adjuvants. At day 8 PI, cells in the lungs and bronco-alveolar lavage (BAL) were stained with D^b^/NP366 and the indicated antibodies. (A), (B), (C) and (D) FACS plots and graphs show percentages of gated tetramer-binding CD8 T cells in respective gates/quadrants or median fluorescence intensities (MFI) for as indicated. Data are representative of three independent experiments. *, **, and *** indicate significance at P<0.1, 0.01 and 0.001 respectively.

We assessed whether adjuvants differed in terms of regulating the differentiation of effector T cells. T cells expressing higher levels of IL-7 receptor (CD127) and lower levels of a terminal differentiation/senescence marker KLRG-1 are associated with greater memory CD8 T cell formation^19^. Of the PLP preparations tested, PLP-CpG reduced CD127 expression, while increasing KLRG-1 expression, such that both PLP-CpG and ADJ+PLP-CpG induced significantly lower frequencies of CD127^HI^/KLRG-1^LO^ CD8 T cells than all other combination PLP adjuvants (Fig 2B). PLP-CpG also drove a strong trend for increased frequencies of CX3CR1^HI^ CD8 T cells (Fig 2B), a marker associated with increased effector differentiation^20^. We quantified transcription factors EOMES and TBET, whicht are known to regulate differentiation of effector CD8 T cells in spleen^21^. Interestingly, while the combination of ADJ+PLP-CpG induced very high levels of transcription factors TBET and EOMES, the combination of ADJ+PLP-GLA had a suppressive effect, leading to lower levels of these transcription factors and lower levels of KI67 expression, a marker of cell proliferation (Fig 2C). Thus, increased expressions of EOMES and TBET were associated with greater terminal differentiation of CD8 T cells in the ADJ+PLP-CpG group.

To determine if combination PLP adjuvants differentially regulated mucosal imprinting of lung CD8 T cells, we analyzed CD69 and CD103 expression. ADJ and PLP-GLA both increased CD69^HI^/CD103^HI^ CD8 T cells in the lungs of vaccinated animals, while PLP-CpG appeared to decrease this mucosal imprinting (Fig. 2D). The combination of ADJ+PLP-GLA led to significantly higher levels of mucosal imprinting in airways and lungs compared with PLP-CpG, ADJ+PLP-CpG, or PLP-GLA. Other markers have been indicated in phenotyping of lung tissue-resident memory cells (TRMs), including CD49a^22^. In our studies, CD49A expression on CD8 T cells appeared to be closely associated with PLP-CpG treatment (Fig 2D), and unlike CD69 and CD103, was expressed to significantly lower levels on CD8 T cells from groups administered ADJ+/-PLP-GLA. To determine the localization of effector CD8 T cells in lungs, we performed vascular staining of T cells shortly before euthanasia. Levels of non-vascular cells were highest in the ADJ and ADJ+PLP-GLA groups, which more closely associated with CD69/CD103 levels than CD49A levels. Thus, ADJ+/-PLP-GLA drives expression of CD103 and CD69, which promote retention of effector CD8 T cells in tissues. By contrast, PLP-CpG appears to dampen the expression of CD103/CD69, leading to the egress of effector T cells from lungs into the vasculature.

TLR agonist-loaded PLPs elicited high frequencies of antigen-specific CD4 T cells in lungs and airways (Fig. 3A). TLR4 and/or 9 agonist-loaded PLP adjuvants induced an accumulation of higher numbers of NP311-specific CD4 T cells in lungs compared to groups that received empty PLPs (Fig 3A). Similar to CD8 T cell responses, the combination of ADJ+PLP-GLA showed a trend for highest levels of antigen-specific responses, however in contrast to NP366-specific CD8 T cell responses, PLP-CpG +/- ADJ also appeared sufficient to induce high frequencies and numbers of NP311-specific CD4 T cells in the respiratory mucosa (Fig 3A).

**Figure 3.**
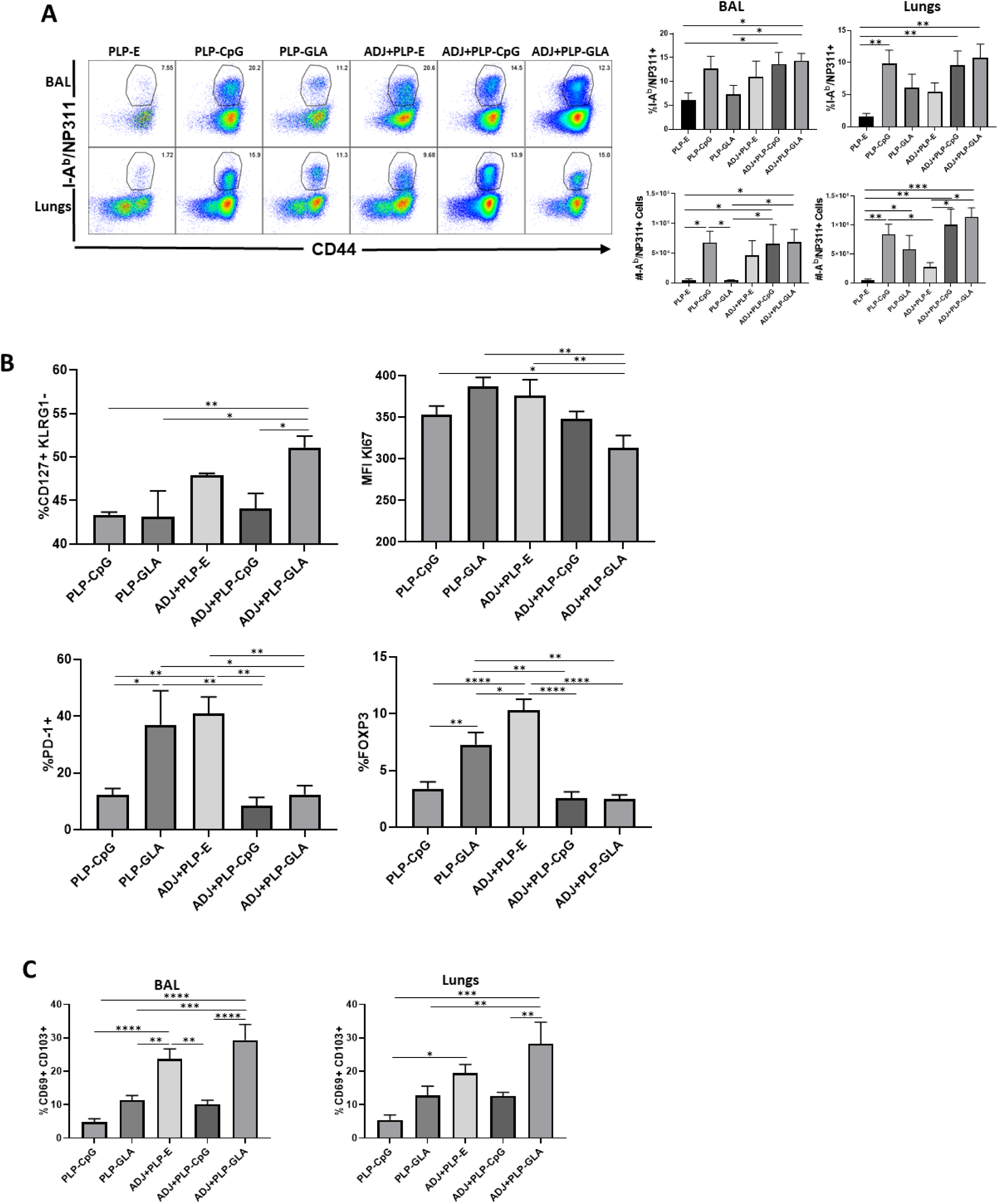
Effector CD4 T cell response to adjuvanted vaccines. At day 8 post vaccination, cells from lungs and BAL were stained with I-A^b^/NP311 tetramers and the indicated antibodies. (A), (B), and (C) FACS plots show percentages of gated tetramer-binding CD4 T cells in respective gates/quadrants or median fluorescence intensities (MFI) for as indicated. Data are representative of three independent experiments. *, **, and *** indicate significance at P<0.1, 0.01 and 0.001 respectively.

Pertaining to the differentiation of effector CD4 T cells, only the combination of ADJ+PLP-GLA significantly increased the frequency of CD127^HI^/KLRG-1^LO^ CD4 T cells, relative to other treatment groups (Fig 3B). The expression of other phenotypic markers (CX3CR1) and transcription factors (TBET/EOMES/IRF4) were not significantly different between treatment groups (supplemental Fig. 1A). However, CD4 T cells isolated from mice that were vaccinated with ADJ+PLP-GLA appeared to be proliferating at significantly lower levels, as measured by lower KI67 staining (Fig 3B). Interestingly, the frequencies of PD-1^HI^ NP311-specific CD4 T cells and FoxP3^HI^ regulatory CD4 T cells were significantly higher in both the PLP-GLA and ADJ+PLP-E groups. Similar to the findings regarding CD8 T cells, ADJ and ADJ+PLP-GLA appeared to drive mucosal imprinting of CD4 T cells, as these groups had significantly higher frequencies of CD69^HI^/CD103^HI^ NP311-specific CD4 T cells. However, CD49A expression and levels of non-vascular T cells were not significantly different between treatment groups (supplemental Fig. 1B).

### Distinct programming of the functionality of CD8 and CD4 T cells by combination PLP adjuvants

Every PLP-based adjuvant elicited strong CD8 Tc1 responses, measured by IFNγ secretion after *ex vivo* NP366 peptide stimulation (Fig 4A). In general, the frequency of IFNγ-secreting CD8 T cells from TLR agonist (CPG/GLA) groups was at least two-folds higher than the ADJ+PLP-E group, indicating TLR4/9 promoted Tc1 programming. Production of IL-17α in CD8 T cells was significantly increased by PLP-GLA-containing adjuvant formulations, yet was barely detectable in PLP-CpG groups, highlighting the contrasting functional programming of TLR4 and TLR9 agonists, respectively (Fig 4A). Though not significant, there was a trend for lower polyfunctionality as measured by frequency of TNFα/IL-2+ among IFNγ+ CD8 T cells in PLP-CpG groups, mirroring the lack of IL-17 production (Fig 4B). Overall, PLP-CpG adjuvants appear to result in strong Tc1 polarization, while vaccination with PLP-GLA resulted in balanced Tc1/Tc17 responses and greater functional diversity. We also evaluated whether adjuvants differed in terms of inducing effector differentiation, as measured by expression of granzyme B. Expression of the effector molecule granzyme B in CD8 T cells was not associated with administration of PLP-CpG, and conversely NP366-specific CD8 T cells in mice receiving PLP-GLA had significantly higher levels for granzyme B, compared to all other adjuvant combinations (Fig 4D).

**Figure 4.**
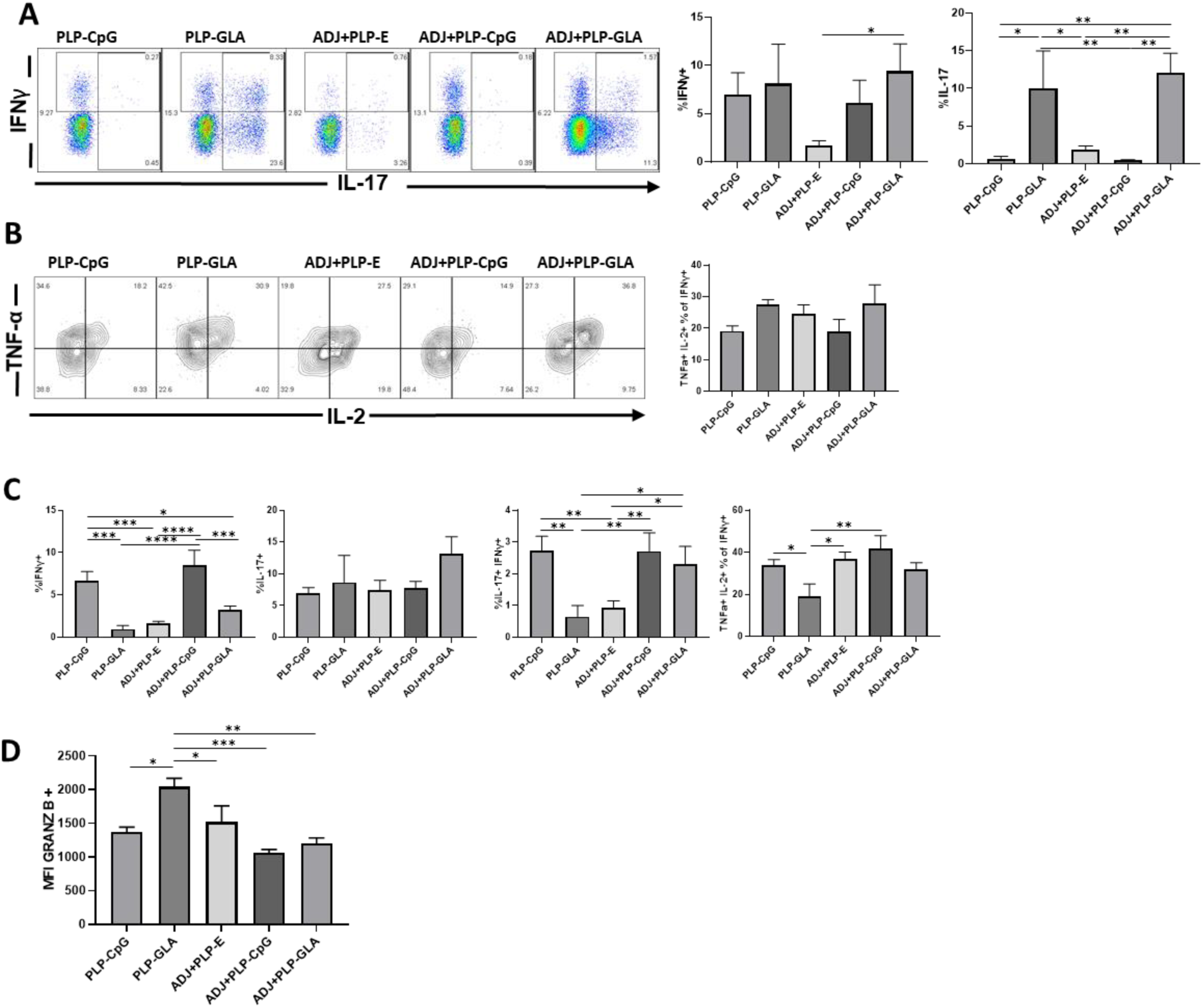
Functional polarization of effector CD8 and CD4 T cells in vaccinated mice. On the 8^th^ day after vaccination, isolated cells were stimulated *ex vivo* with NP366 or NP311 peptides for 5 hrs. The percentages of NP366-specific CD8 T cells or NP311-specific CD4 T cells that produced IFN-γ, IL-17, TNF-α, and IL-2 were quantified by intracellular cytokine staining. FACS plots and graphs show the percentages of cytokine-producing cells among the gated CD8 T cells. Graphs are gated on IFN-γ-producing CD8 T cells. Graphs show the percentages of cytokine-producing cells among CD4 T cells as above (C). Data are representative of three independent experiments. *, **, and *** indicate significance at P<0.1, 0.01 and 0.001 respectively.

In stark contrast to CD8 T cell functionality, PLP-CpG tended to drive potent and balanced Th1/Th17 responses in CD4 T cells, compared with PLP-GLA only administration. PLP-CpG +/- ADJ groups had significantly higher frequencies of IFNγ+ CD4 T cells, while all combination PLP adjuvant groups exhibited similarly high frequencies of IL-17 secretion (Fig 4B). Further, the PLP-GLA group had lower frequencies of polyfunctional triple cytokine-producing (IFNγ, TNF, and IL-2) CD4 cells than the ADJ+PLP-GLA and ADJ +/- PLP-CpG groups (Fig 4B). When the functional polarization of CD4 and CD8 T cells is taken together, ADJ+PLP-GLA seems to induce the most balanced and potent Tc1/Tc17 and Th1/Th17 program. These results also demonstrate that the type of TLR agonist conjugated to PLPs can have disparate programming effects on CD4 and CD8 T cell functionality during the effector phase of vaccination.

### PLP adjuvants affect the magnitude and functionality of CD4 and CD8 T cell memory

While all combination PLP adjuvants elicited durable antigen (NP366)-specific CD8 T cell memory at D100 post boost, the ADJ+PLP-GLA group had significantly higher frequencies and/or total numbers of memory CD8 T cells in both airways and lungs than other groups (Fig 5A). Phenotypically, memory CD8 T cells from all groups were relatively homogeneous at this time point. There were no substantial differences in CD49a, CD62L, CD69, CD103, CD127, CXCR3, CX3CR1, or KLRG1+ frequencies among NP366-specific memory CD8 T cells. However, the numbers of non-vascular parenchymal tissue-resident memory CD8 T cells in lungs of ADJ+PLP-GLA mice were significantly higher, as compared to other adjuvant groups (Fig 5A)

**Figure 5.**
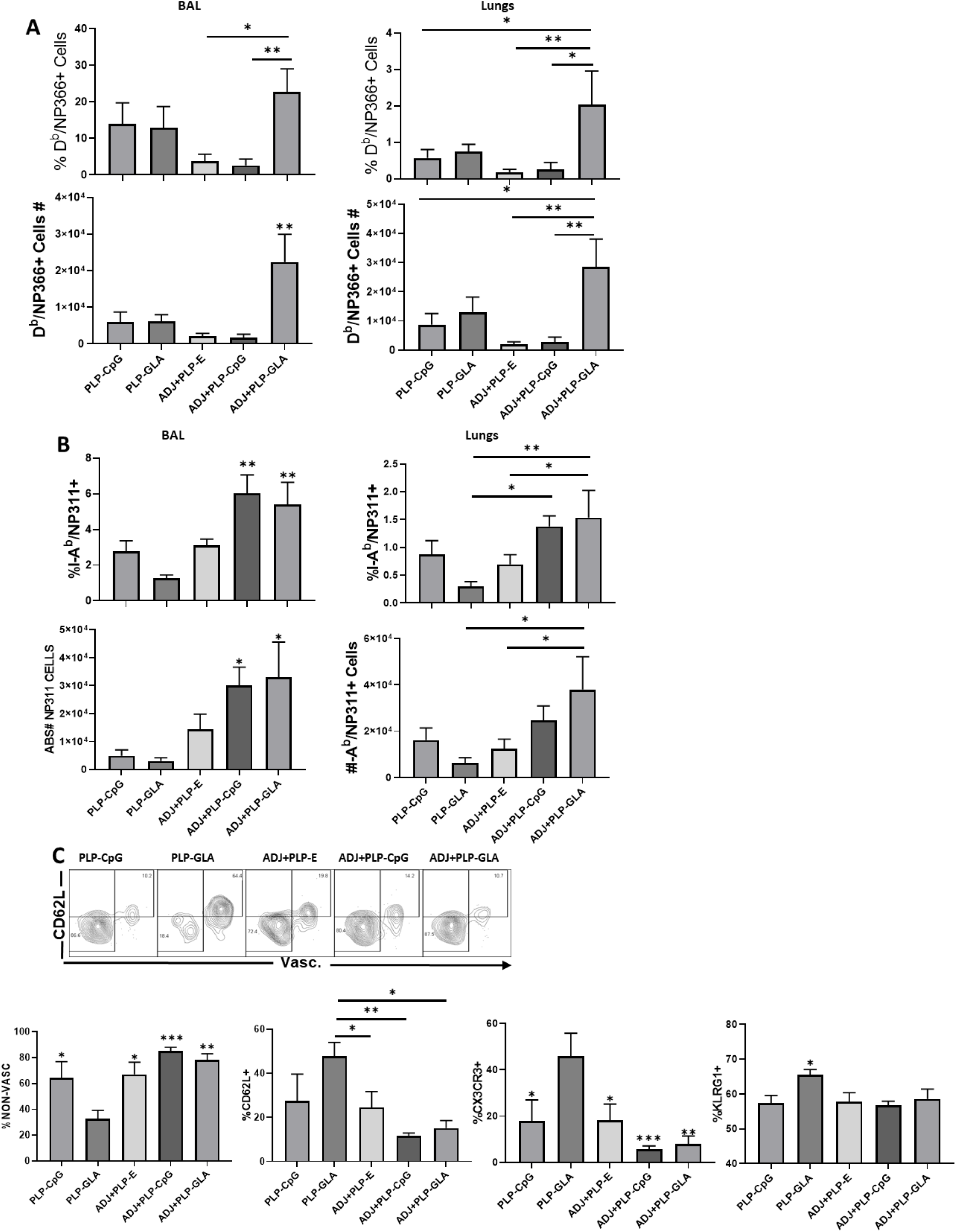
Mucosal CD8 and CD4 T cell memory in vaccinated mice. At 100 days after vaccination, to stain vascular cells, mice were injected intravenously with PE-labeled anti-CD4.2 antibody. Cells from BAL and lungs were stained with D^b^/NP366 tetramers, I-A^b^/NP311 tetramers. Percentages and total numbers of NP366-specific CD8 T cells or NP311-specific CD4 T cells in BAL or lungs (A, B respectively). Plots in (C) are gated on NP311-specific T cells. Data are representative of three independent experiments. *, **, and *** indicate significance at *P*<0.1, and 0.001 respectively.

ADJ+PLP-CpG and ADJ+PLP-GLA groups had significantly higher frequencies and numbers of antigen (NP311)-specific memory CD4 T cells in airways than other groups and both groups had significantly higher numbers in airways and/or lungs than the PLP-GLA group (Fig 5B). Among these NP311-specific CD4 T cells, the PLP-GLA group had a significantly lower proportion of cells in the lung parenchyma (non-vascular) and a significantly higher frequency of CD62L expression (among vascular memory T cells) than other groups (Fig 5B). Interestingly, at this memory time point, CD4 T cells from the PLP-GLA group also expressed effector markers CX3CR1 and KLRG1 at significantly higher levels than other groups.

Functionally, NP366-specific IFNγ+ memory CD8s were not substantially different between groups (Fig 6A), though similar to D8, the percentages of IFNγ+ CD8 T cells in TLR4 or 9 agonist groups were generally higher than ADJ+PLP-E, and the ADJ+PLP-GLA group had the highest IFNγ+ frequencies. Interestingly, while both ADJ+PLP-GLA or ADJ+PLP+CpG groups had strong Tc1 responses, the degree of polyfunctionality (ability to also produce TNFα and IL-2) among IFNγ producers in the ADJ+PLP-GLA group was significantly higher than the ADJ+PLP-CpG group. IL-17-producing CD8 T cells in the ADJ+PLP-GLA group were significantly higher than in all other groups (Fig 6A). While Tc17 responses in all groups contracted from D8 to D100, this contraction seemed to occur to a lesser degree in the ADJ+PLP-GLA group than the ADJ-GLA group; on D8, the numbers and frequencies of NP366-specific CD8 T cells, and frequencies of Tc17 cells were not different between these two groups, but at D100, Tc17 frequencies were significantly higher in ADJ+PLP-GLA than the PLP-GLA group.

**Figure 6.**
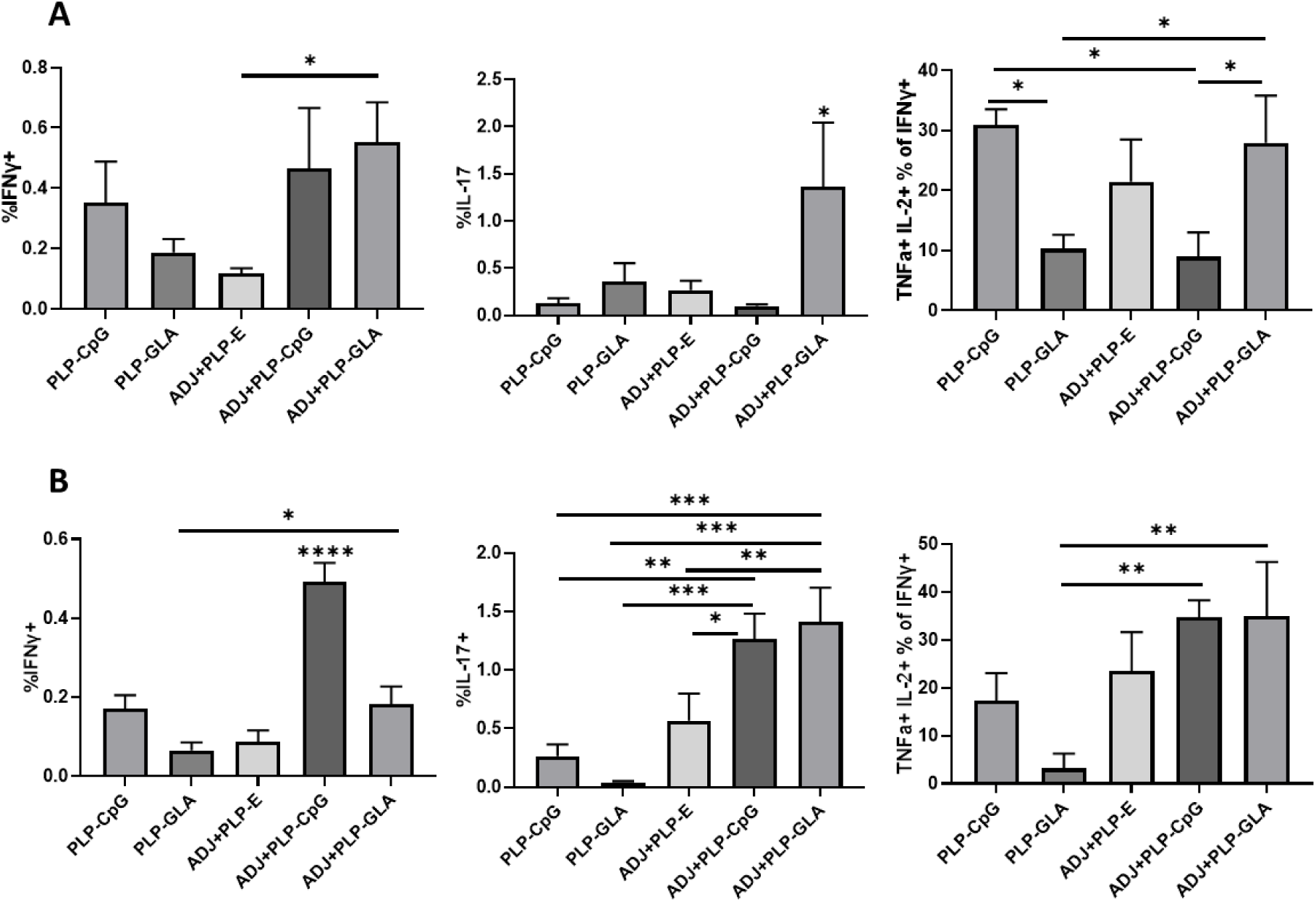
Functional polarization of memory CD8 and CD4 T cells in vaccinated mice. Percentages of IFN-γ, IL-17, or TNF-α and IL-2 producing cells among NP366-specific CD8 T cells (A) or NP311-specific CD4 T cells (B). Data are representative of three independent experiments. *, **, and *** indicate significance at *P*<0.1, 0.01 and 0.001 respectively.

All groups had robust IFNγ+ memory CD4 T cells at D100, however, ADJ+PLP-CpG had significantly higher frequencies of NP311-specific memory T cells (Fig 6B). Further, NP311-specific memory CD4 T cells in the ADJ+PLP-GLA and ADJ+PLP-CpG groups appeared to have greater overall functionality, as IFNγ+ cells also secreted IL-2 and TNFα to greater levels, and both groups had significantly higher frequencies of IL-17-producing cells (Fig 6B). When the functionality of both memory CD4 and CD8 cells are taken together, we observed a similar trend, which is that ADJ+PLP-GLA appeared to induce the most robust and balanced Tc1/Th1 and Th17/Tc17 responses.

### ADJ+PLP-GLA induces durable and potent immunity to influenza virus challenge

Cohorts of vaccinated animals were challenged with a lethal dose of H1N1 PR8 influenza virus at D101 post boost. At D6 post challenge, we quantified recall T cell responses and viral titers in the lungs. While vaccination with all PLP formulations resulted in at least an approximately 2 Log10 reduction in lung viral titers, the ADJ+PLP-GLA vaccine conferred the highest degree of viral control, reducing viral titers by nearly 5 Log10 PFU/gram of lung compared to mock (saline) vaccinated mice (Fig 7A). 4/5 animals in the ADJ+PLP-GLA group had no detectable infectious virus by plaque assay even in undiluted lung homogenate, indicating nearly complete control of lung viral replication by this D6 time point.

**Figure 7.**
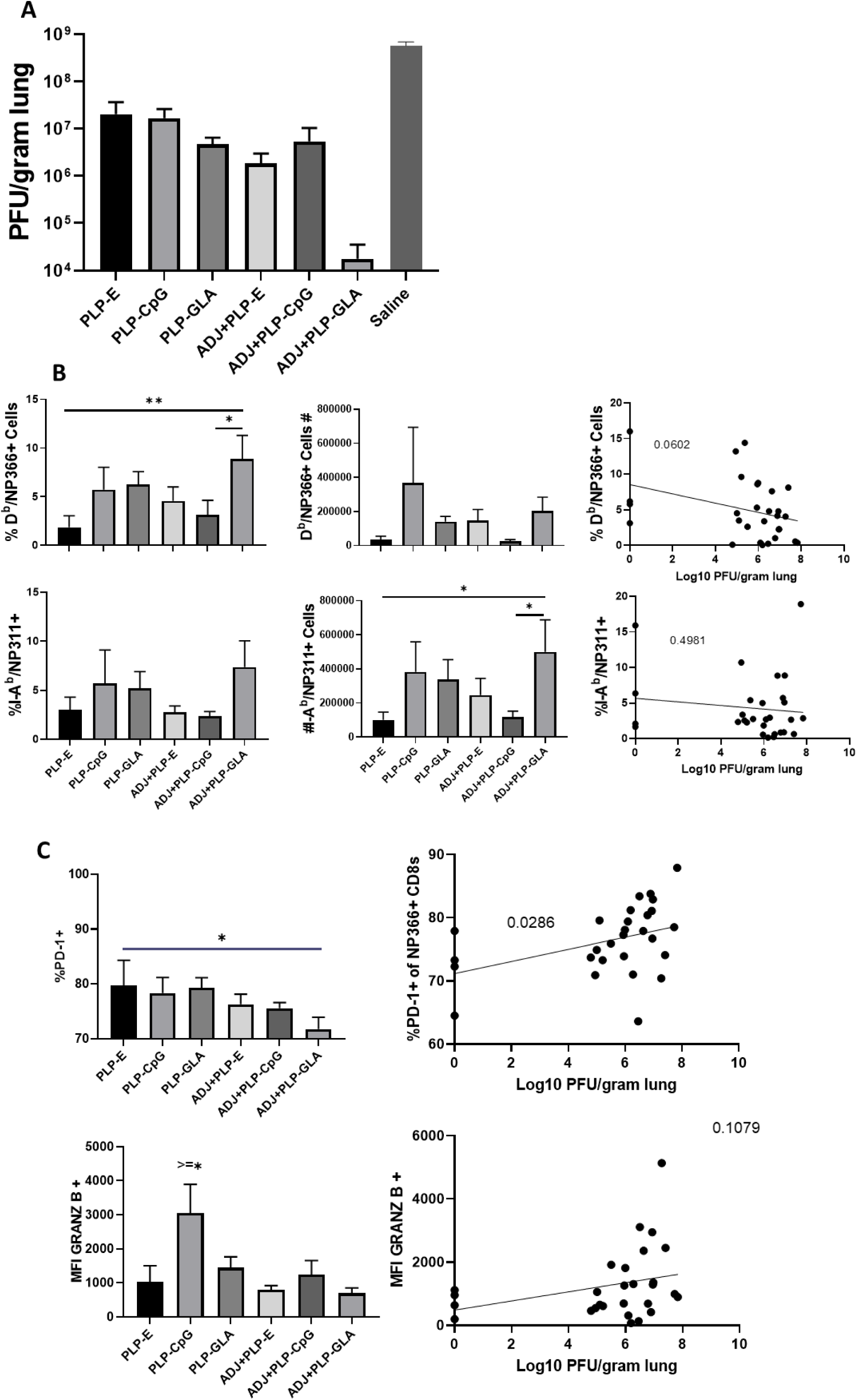
Vaccine-induced protective immunity influenza virus. 101 days after vaccination with PLP adjuvants, mice were challenged with H1N1/PR8 strain of influenza A virus. Viral titers were quantified in the lungs on D6 after challenge (A). Numbers of NP366-specific CD8 T cells and NP311-specific CD4 T cells in lungs (B). Plots are gated on NP366 CD8 T cells (C). Data are representative of two independent experiments. *, **, and *** indicate significance at *P*<0.1, 0.01, and 0.001 respectively. Linear regression curves were plotted for data from individual mice for the indicated cell frequency plotted against its Log10 viral titer value.

Interestingly, total numbers and frequencies of recall antigen-specific CD4 and CD8 T cells did not vary significantly between vaccinated groups at this time point (Fig 7B). However, there was a nearly significant relationship between frequencies of NP366-specific CD8s and viral lung titers (Fig 7B), whereas frequencies or numbers of CD4 T cells did not exhibit such a strong trend. Additionally, there were no noteworthy phenotypic or transcriptional differences in CD8 T cells in terms of CD49a, CD62L, CD69, CD103, CD127, CXCR3, CX3CR1, KLRG1, TBET, EOMES, and IRF4 expression. However, we noticed a strong trend for reduced PD-1 expression on recall CD8 T cells in ADJ containing groups, particularly for ADJ+PLP-GLA, and there was a strong negative correlation between PD-1 expression levels and viral lung titers (Fig 7C). Further, granzme B levels were significantly upregulated in recall CD8 T cells in the PLP-CpG group, and there was a negative trend for granzyme levels and viral titer. Increased PD-1 and granzyme B expression in CD8 T cells likely reflects ongoing antigenic stimulation and CTL function in mice, asscociated with delayed viral clearance. Finally, we did not observe any strong or clear trends between the aforementioned markers on recall CD4 T cells and viral control.

### Recall T cell function intimately associates with viral control

All PLP groups showed robust IFNγ+ CD8 T-cell recall responses measured by *ex vivo* NP366 peptide stimulation (Fig 8A). There was a clear and strong association between the frequency of IFNγ expression in CD8s and the degree of viral control in lungs (Fig 8A). ADJ seemed to drive increased functionality among IFNγ-producing recall CD8 T cells (Fig 8A). With the exception of ADJ+PLP-CpG, both ADJ and PLP-GLA groups had significantly increased frequencies of IL-17 expression during virus-induced CD8 T cell recall, with the ADJ+PLP-GLA group having significantly higher levels than all other groups (Fig 8A). Indeed, ADJ+PLP-GLA had the greatest Tc17 recall response and the lowest lung viral titers, and overall there was an extremely close association between IL-17 expression in CD8s and viral control (Fig 8A), such that even removing the ADJ+PLP-GLA animals that had no detectable lung titer from the correlation analysis still resulted in a significant association (supplemental Fig 2).

**Figure 8.**
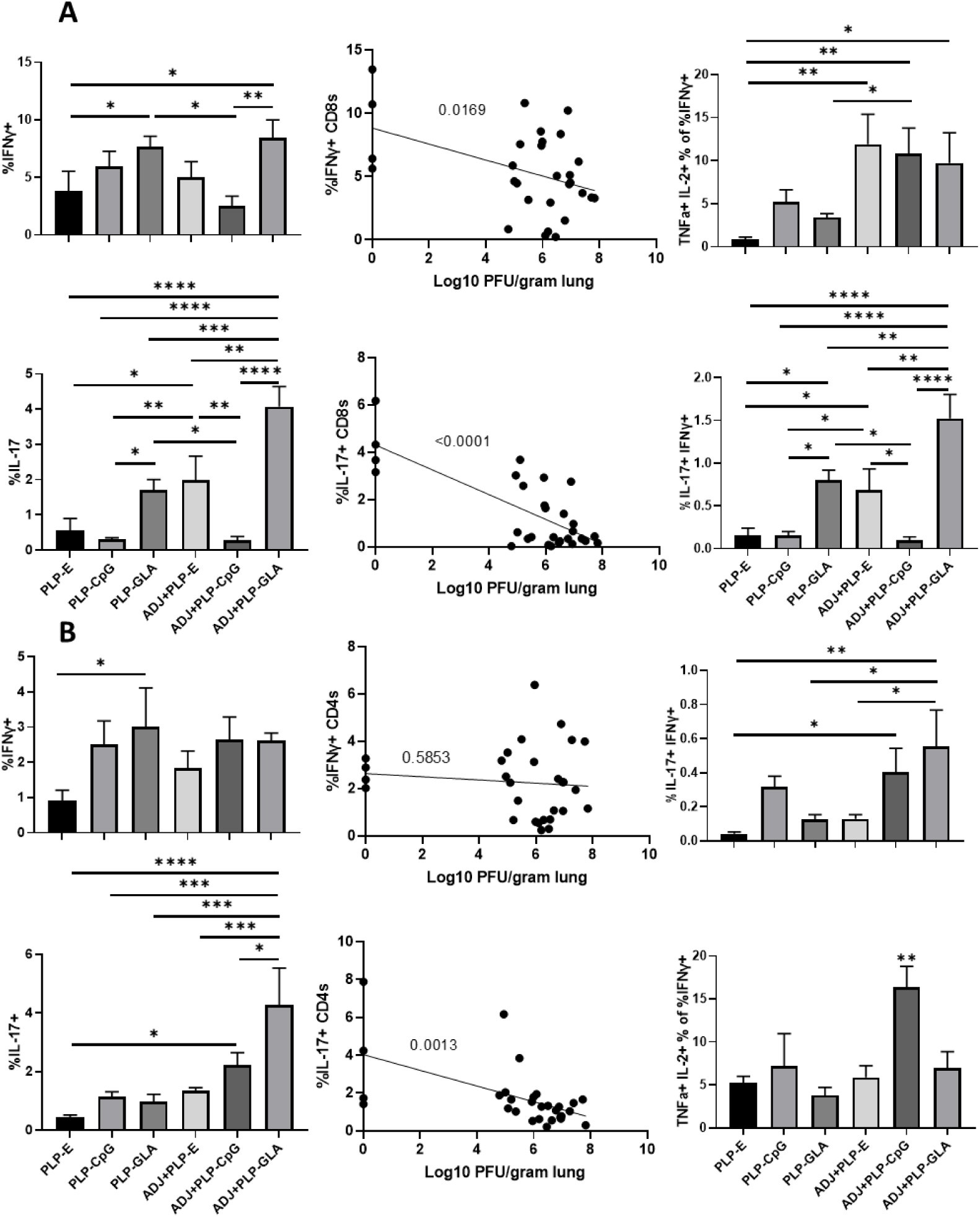
Functional polarization of recall CD8 and CD4 T cells. 101 days after vaccination with PLP adjuvants, mice were challenged with H1N1/PR8 strain of influenza A virus. Isolated cells were stimulated *ex vivo* with NP366 or NP311 peptides for 5 hrs. The percentages of NP366-specific CD8 T cells or NP311-specific CD4 T cells that produced IFN-γ, IL-17, TNF-α and IL-2 were quantified by intracellular cytokine staining. Graphs show the percentages of cytokine-producing cells among the gated CD8 T cells (A). Graphs show the percentages of cytokine-producing cells among CD4 T cells as above (B). Data are representative of two independent experiments. *, **, and *** indicate significance at P<0.1, 0.01, and 0.001 respectively. Linear regression curves were plotted for data from individual mice for the indicated cell frequency plotted against its Log10 viral titer value.

For CD4 Th1 recall responses, while all groups had robust IFNγ+ CD4 T cell responses, there were no noteworthy differences between groups and there did not appear to be a strong association with viral control (Fig 8B). On the other hand, the ADJ+PLP-GLA group had a significantly higher recall Tc17 response than other groups, and frequencies of IL-17+ expression in CD4s significantly correlated with reduced viral lung titer (Fig 8B). Overall, the robust and balanced Tc1/Th1 and Th17/Tc17 functional responses elicited by the ADJ+PLP-GLA vaccine appeared to result in highly efficient viral control.

## DISCUSSION

Programming potent and durable B and T cell memory is the goal of vaccination programs. Live viral vaccines such as the Yellow fever vaccine (YFV) and the small pox vaccine engender immunity that lasts for decades after vaccination^23^. Studies of YFV suggest that engagement of multiple innate immune receptors early in the infection may be key to the programming of long-lived immunological memory^24^. This paradigm has triggered a multitude of investigations to explore the possibility of using TLR agonists as adjuvants to program durable immunological memory. One of the downsides of using TLR agonists, such as CpG, in soluble form is their diffusion from vaccination site, leading to systemic toxicity in vaccinees^25^. To circumvent this problem and mimic the biophysical attributes of pathogens and their interactions with pattern recognition receptors, we engineered biodegradable PLGA microparticles (i.e. PLPs) that were loaded with optimized densities of TLR agonists CpG and GLA^15^. In this manuscript, we document for the first time that mucosal delivery of CpG-or GLA-loaded PLPs elicit unexpectedly potent mucosally imprinted antigen-specific CD4 and CD8 T cell responses in the respiratory tract. Interestingly, we find that PLP-CpG and PLP-GLA stimulate disparate transcriptional programs that evoke distinct phenotypic and functional differentiation of antigen-specific T cells. Further, we show that the combination of PLP-GLA, but not PLP-CpG, with the nanoemulsion adjuvant ADJ synergistically augmented the magnitude of lung-resident T cell memory and protective immunity to a lethal influenza virus infection. These studies highlight how the mode of TLR agonist presentation can be leveraged to achieve enhanced immunogenicity without toxicity, and how this feature can be combined with the antigen presenting properties of a nano-emulsion adjuvant to program effective T cell-based protective immunity in the respiratory tract.

*In vitro* studies show that soluble/PLP-CpG, but not soluble/PLP-MPLA, triggers IFNβ production by BMDCs. Conversely, only soluble/PLP-GLA, but not CpG, (supplementary Fig. 3) stimulate IL-1β production when combined with ADJ. Despite this difference in IFNβ induction and inflammasome activation, both PLP-CpG and PLP-GLA elicited strong, yet comparable levels of effector CD8 and CD4 T cell responses in vivo. This data suggests that (1) lack of IFNβ induction in vitro does not predict failure by an adjuvant to induce T cell responses in vivo; (2) PLP-CpG and PLP-GLA likely stimulate different arrays of cytokines in DCs in vivo; (3) PLP-CpG and PLP-GLA might engage different pathways to stimulate T cell activation and expansion in the respiratory tract. Consistent with this idea, it is noteworthy that PLP-CpG and PLP-GLA differ when the differentiation of T cells are compared. In comparison to PLP-GLA, PLP-CpG tended to drive T cells towards terminal differentiation, based on elevated expressions of KLRG-1, CX3CR1, Tbet, and EOMES. CpG is known to induce high levels of IL-12^26^, and it is likely that higher induction of IL-12 in DCs by PLP-CpG underlies greater levels of Tbet and terminal differentiation of effector T cells in PLP-CpG-vaccinated mice. Pertaining to functional polarization of T cells, PLP-CpG elicited primarily Tc1 CD8 T cells and Th1 CD4 T cells secreting IFNγ, but only a fraction of CD4 T cells also secreted IL-17α. In contrast, PLP-GLA promotes functionally broad CD8 and CD4 T cell responses that secreted IFNγ and/or IL-17α. The differential functional polarization of T cells in PLP-CpG and PLP-GLA-vaccinated mice might be linked to disparate levels of IL-12 and inflammasome activation, respectively^15,26,27^.

Total numbers of influenza lung tissue-resident memory CD8 T cells correlate with protection from rechallenge^8-10^. We have previously reported that ADJ, a carbomer-containing nanoemulsion adjuvant induces tissue-resident memory T cells in lungs and protects against pathogenic influenza A virus infection^13^. In the current study, we explored whether combining ADJ with PLP-CpG or PLP-GLA augmented the adjuvanticity of ADJ and increased the numbers of lung-resident memory CD8 T cells. PLP-CpG, PLP-GLA, and ADJ alone did not significantly differ in terms of the numbers of lung TRM CD8 T cells. Although ADJ suppresses PLP-CpG-induced IFNβ production in vitro, ADJ did not adversely affect CD8 or CD4 T cell responses to PLP-CpG. The numbers of effector CD8 T cells (at the peak of the response) or memory CD8 T cells were not significantly different in PLP-CpG versus ADJ+PLP-CpG groups. Thus, combining ADJ with PLP-CpG did not alter the development of memory CD8 T cells. In striking contrast, combining ADJ with PLP-GLA generally enhanced the number of effector and memory CD8/CD4 T cells, as compared to ADJ or PLP-GLA alone; ADJ+PLP-GLA induced the largest cohort of lung- and airway-resident memory T cells, as compared to all other groups. This in turn correlated strongly with the most effective protective immunity against pathogenic influenza virus infection. As discussed above, PLP-CpG with or without ADJ promotes high level of expression of TBET leading to a greater degree of differentiation of KLRG-1^HI^/CX3CR1^HI^ CD8 T cells. By contrast, the combination of ADJ+PLP-GLA maximizes effector/memory mucosal NP366-specific CD8 T cells and TRM frequencies, and downregulates the levels of KLRG-1, CX3CR1, Granzyme B, TBET, EOMES, and Ki67 ^28,29^. Furthermore, engaging the RORγ/Tc17/Th17 differentiation program by ADJ+PLP-GLA might enhance differentiation of long-lived stem cell-like memory T cells^30^. Nonetheless, our results are consistent with the axiom that TRMs arise from less differentiated effector cells, and terminally differentiated effector cells have diminished capacity to develop into TRMs^31,32^. Collectively, our findings support the rationale and feasibility of combining adjuvants to mitigate terminal differentiation of effectors and enhance the development of TRMs.

Memory T cell-dependent protection against influenza virus is determined by the number of memory CD8 T cells in airways and lung parenchyma^8-10,33^. Consistent with these reports, we find that ADJ+PLP-GLA mice contained significantly greater numbers of TRMs and memory T cells in airways, which correlated strongly with the most effective protection against pathogenic influenza infection. Additionally, we find a strong correlative link between recall CD4/CD8 T cell functionality and reduction of influenza virus lung titers, specifically in Tc1/Tc17 and Th1/Th17 responses. The role of IFNγ, and its production in recall lung CD8 T cell responses is established in controlling influenza viral lung replication^34-37^. While we also observed a strong correlation between IFNγ expression in lung CD8 T cells and reduction in viral titers (Fig 8A), this association did not apply for lung CD4 T cells (Fig 8B), indicating that perhaps Tc1 cells are more important than Th1 cells in controlling influenza virus. Interestingly, production of IL-17α from both CD8 and CD4 T cells was closely associated with reduction in lung viral lung titers in this study (Fig 8A, 8B). IL-17α-producing T cells are well known to be involved in fungal immunity^38,39^, however the role of Tc17/Th17 responses in control of influenza infection is only recently emerging. Lung Tc17 cells appear to be a distinct subset of CD8 T cells, compared with Tc1 or Tem cells, and are associated with protection from rechallenge^40,41^. Influenza-specific lung Th17 have also been implicated in protection from influenza^42,43^. Overall, more mechanistic studies are required to carefully dissect and correlate the roles of individual functional responses in vaccine-induced lung T cells to influenza viral control.

While antibody-mediated protection against influenza virus is type and subtype specific, memory T cells that recognize conserved epitopes in the internal proteins, such as nucleoprotein, provide heterosubtypic immunity to influenza A virus^44,45^. Hence, there is an impetus to identify strategies to elicit T cell immunity in the lungs towards an universal influenza vaccine^44^. However, there are no FDA-approved mucosal adjuvants that are known to elicit protective T cell immunity in the respiratory mucosa. In this manuscript, we have explored novel ways of presenting immune components to the immune system to maximize antigen presentation to T cells and evoke innate immune responses that program a strong and enduring mucosal T cell response in the respiratory tract. Specifically, we have identified a novel vaccine formulation consisting of influenza virus NP, TLR-loaded PLPs, and a nanoemulsion adjuvant, that elicits robust mucosally-imprinted T cell memory in the respiratory tract and affords effective protective immunity to a pathogenic influenza virus infection in mice. These findings have significant implications in the development of T cell-based universal influenza vaccines for humans and animals.

## Methods

### Experimental animals

6-12 week-old C57BL/6 (B6) were purchased from Jackson Laboratory or from restricted-access SPF mouse breeding colonies at the University of Wisconsin-Madison Biotron Laboratory. All mice were housed in specific-pathogen-free conditions in the animal facilities at the University of Wisconsin-Madison (Madison, WI). Knockout mice used for *in vitro* studies were bred in the lab of Dmitry Shayakhmetov at Emory University.

### Ethics statement

These studies were carried out in strict accordance with recommendations set forth in the National Institutes of Health Guide for the Care and Use of Laboratory Animals. All animals and animal facilities were under the control of the School of Veterinary Medicine with oversight from the University of Wisconsin Research Animal Resource Center. The protocol was approved by the University of Wisconsin Animal Care and Use Committee (Protocol number V005308). The animal committee mandates that institutions and individuals using animals for research, teaching, and/or testing much acknowledge and accept both legal and ethical responsibility for the animals under their care, as specified in the Animal Welfare Act (AWA) and associated Animal Welfare Regulations (AWRs) and Public Health Service (PHS) Policy.

### Cells and viruses

Murine bone marrow-derived dendritic cells (BMDCs) were differentiated from bone marrow isolated from C57BL/6, AIM2^-/-^, NLRP3^-/-^, AIM2/NLRP3^-/-^, or ASC^-/-^ mice. The bone marrow was processed into single cell suspensions and treated with RBC lysis buffer. Cells were then plated into Petri dishes and cultured in BMDC differentiation medium (RPMI, 10% FBS, 1% penicillin-streptomycin, 1x sodium pyruvate, 1x β-mercaptethanol) supplemented with GM-CSF (Peprotech, Rocky Hill, NJ). Media was refreshed on day 2, day 4, and day 6. On day 7, BMDCs were harvested and replated for experiments.

Madin-Darby canine kidney (MDCK) cells were obtained from ATCC (ATCC; Manassas, VA, USA) and propagated in growth media containing Modified Eagle’s Medium (MEM) with 10% fetal bovine serum (FBS; Hyclone, Logan, UT), 2 mM L-glutamine, 1.5 g/l sodium bicarbonate, non-essential amino acids, 100 U/ml of penicillin, 100 μg/ml of streptomycin (flu media), and incubated at 37°C in 5% CO2. Reverse genetics-derived influenza virus strain A/PR/8/34 H1N1 (PR8) were propagated in MDCK) cells, and viral titers were determined by plaque-forming assay^46,47^. Briefly, MDCK cells grown to 90% confluency were infected with serial dilutions of influenza virus samples, and incubated for 1hr while periodically shaking under growth conditions. Cells were then washed with PBS and incubated in flu media containing 1% SeaPlaque agarose (Lonza, Basel, Switzerland). After 48hr incubation, cells were fixed in 10% neutral buffered formalin (NBF), agarose plugs were removed, and distinct plaques were counted at a given dilution to determine the plaque forming units (PFU) of virus per sample.

### Viral challenge

For PR8 challenge studies, mice were inoculated with 500 PFU of PR8 by intranasal (IN) instillation in 50ul PBS applied to the nares under isoflurane anesthesia, and were humanely euthanized at 6 days post infection. Lung tissues for viral titration (left lobe) were frozen at -80°C.

### Vaccines and vaccinations

PR8 nucleoprotein (NP) was purchased from Sino Biological Inc (Beijing, China). CpG ODN 1826 (CpG) oligonucleotide adjuvant was purchased from InivivoGen (San Diego, CA). The synthetic monophosphoryl lipid A adjuvant, Glucopyranosyl Lipid Adjuvant (GLA) was purchased from Avanti Polar Lipids, Inc. (Alabaster, AL). PLPs were synthesized by the double emulsion method. Briefly, PLGA was dissolved in dichloromethane in the presence or absence of GLA adjuvant (10 μg GLA/mg PLGA). DI H2O was added and the mixture was homogenized to create the first emulsion. 1% PVA was then added and the mixture was homogenized to create the second emulsion. Excess dichloromethane was removed by solvent evaporation and particles were washed with DI H2O by centrifugation. Following lyophilization, branched PEI was conjugated to the PLP surface by reaction with EDC and sulfo-NHS (Thermo Fisher Scientific, Bedford, MA). PLPs were washed again sequentially with 1 mM NaCl and DI H2O. CpG adjuvant was loaded onto PLPs (10 μg CpG/mg PLGA) without GLA in sodium phosphate buffer (pH = 6.5) overnight at 4C. Particle size and zeta potential at pH 7.4 was measured using a Malvern Zetasizer. All vaccinations were given via IN instillation under isoflurane anesthesia in 50μl saline with 10μg NP adjuvant as follows: 10% ADJ (ADJ) +/-; 1mg PLGA (PLP-E); 1mg PLGA loaded with 10μg CpG (PLP-CpG); 1mg PLGA loaded with 10 μg GLA (PLP-GLA); 10% ADJ.

### BMDC activation and proliferation

Murine BMDCs were plated in 96-well plates (300,000 cells/well). BMDCs were incubated with ADJ (1%) and/or PLP adjuvants (50 μg PLGA/mL). After 24 hours, supernatants were collected. IFN-β, IL-1β, and IL-18, were measured by ELISA (BioTechne, Minneapolis, MN). Cells were then incubated with CellTiter 96 Aqueous One Solution Proliferation Solution for 1 hour (Promega, Fitchburg, WI). Absorbance of the solution was then read at 490 nm. Measurements were normalized to untreated cells at the same timepoint of incubation.

### Flow cytometry

For indicated studies, vascular staining of T-cells was performed by IV injection of fluorochrome-labeled CD45.2 three minutes prior to animal euthanasia and necropsy. Single-cell suspensions from spleen, lung, and bronchoalveolar lavage were prepared using standard techniques. Prior to antibody staining, cells were stained for viability with Fixable Viability 780 (eBiosciences, San Diego, CA) according to manufacturer’s instructions. Fluorochrome-labeled antibodies against the cell-surface antigens, Ly5.2 (CD45.2), CD4, CD8a, CD44, CD62L, KLRG-1, CD127, CD103, CD69, CD49A, CD127, CXCR3, CX3CR1, and intracellular antigens IFN-γ, TNF-α, IL-2, IL-17, TBET, EOMES, IRF-4, and granzyme B antibodies were purchased from BD Biosciences (San Jose, CA), Biolegend (San Diego, CA), eBioscience (San Diego, CA), Invitrogen (Grand Island, NY), or Tonbo Biosciences. Fluorochrome-conjugated I-A^b^ and H2-D^b^ tetramers bearing influenza nucleoprotein peptides, QVYSLIRPNENPAHK (NP311) and ASNENMETM (NP366) respectively, were kindly provided by the NIH Tetramer Core Facility (Emory University, Atlanta, GA). For class-II tetramer NP311, cells were incubate at 37C for 90 min. For class-I tetramers, cells were incubated with tetramer and antibodies for 60 minutes on ice in the dark. Stained cells were fixed with 2% paraformaldehyde in PBS for 20 minutes, then transferred to FACS buffer. All samples were acquired on a LSRFortessa (BD Biosciences) analytical flow cytometer. Data were analyzed with FlowJo software (TreeStar, Ashland, OR).

### Intracellular cytokine stimulation

For intracellular cytokine staining, 1×10^6^ cells were plated on flat-bottom tissue-culture-treated 96-well plates. Cells were stimulated for 5 hours at 37C in the presence of human recombinant IL-2 (10 U/well), and brefeldin A (1 μl/ml, GolgiPlug, BD Biosciences), with one of the following peptides: NP366, NP311 (thinkpeptides^®^, ProImmune Ltd. Oxford, UK) at 0.1ug/ml, or without peptide. After stimulation, cells were stained for surface markers, and then processed with Cytofix/Cytoperm kit (BD Biosciences, Franklin Lakes, NJ).

### Statistical analyses

Statistical analyses were performed using GraphPad software (La Jolla, CA). All comparisons were made using using an ordinary one-way ANOVAtest with Tukey corrected multiple comparisons where p<0.05 = *, p<0.005 = **, p<0.0005 = ***, etc. were considered significantly different among groups. Viral titres were log transformed prior to analysis.

